# Frequencies of circulating Th1-biased T follicular helper cells in acute HIV-1 infection correlate with the development of HIV-specific antibody responses and lower set point viral load

**DOI:** 10.1101/304329

**Authors:** Omolara Baiyegunhi, Bongiwe Ndlovu, Funsho Ogunshola, Nasreen Ismail, Bruce D. Walker, Thumbi Ndung’u, Zaza M. Ndhlovu

**Affiliations:** HIV Pathogenesis Programme, Doris Duke Medical Research Institute, Nelson R. Mandela School of Medicine, University of KwaZulu-Natal, Durban, South Africa.; Ragon Institute of Massachusetts General Hospital, Massachusetts Institute of Technology, and Harvard University, Cambridge, MA, USA.; Howard Hughes Medical Institute, Chevy Chase, Maryland, USA.; Africa Health Research Institute (AHRI), Nelson R. Mandela School of Medicine, University of KwaZulu-Natal, Durban, South Africa.; Max Planck Institute for Infection Biology, Berlin, Germany.

**Keywords:** HIV, T follicular helper cells, non-neutralizing antibodies, Gag p24 IgG

## Abstract

Despite decades of focused research, the field has yet to develop a prophylactic vaccine. In the RV144 vaccine trial, non-neutralizing antibody responses were identified as a correlate for prevention of HIV acquisition. However, factors that predict the development of such antibodies are not fully elucidated. We sought to define the contribution of circulating T follicular helper (cTfh) cell subsets to the development of non-neutralizing antibodies in HIV-1 clade C infection. Study participants were recruited from an acute HIV-1 clade C infection cohort. Plasma anti-gp41, -gp120, -p24 and -p17 antibodies were screened using a customized multivariate Luminex assay. Phenotypic and functional characterization of cTfh were performed using HLA class II tetramers and intracellular cytokine staining. In this study, we found that acute HIV-1 clade C infection skewed differentiation of functional cTfh subsets towards increased Tfh1 (p=0.02) and Tfh2 (p<0.0001) subsets, with a concomitant decrease in overall Tfh1-17 (that shares both Tfh1 and Tfh17 properties) (p=0.01) and Tfh17 subsets (p<0.0001) compared to HIV negative subjects. Interestingly, the frequencies of Tfh1 during acute infection (5.0-8.0 weeks post-infection) correlated negatively with set point viral load (p=0.03, r=-60) and were predictive of p24-specific plasma IgG titers at one year of infection (p=0.003, r=0.85). Taken together, our results suggest that circulating the Tfh1 subset plays an important role in the development of anti-HIV antibody responses and contributes to HIV suppression during acute HIV-1 infection. These results have implications for vaccine studies aimed at inducing long lasting anti-HIV antibody responses.

**Importance:** The HIV epidemic in southern Africa accounts for almost half of the global HIV burden with HIV-1 clade C being the predominant strain. It is therefore important to define immune correlates of clade C HIV control that might have implications for vaccine design in this region. T follicular helper (Tfh) cells are critical for the development of HIV-specific antibody responses and could play a role in viral control. Here we showed that the early induction of circulating Tfh1 cells during acute infection correlated positively with the magnitude of p24-specific IgG and was associated with lower set point viral load. This study highlights a key Tfh cell subset that could limit HIV replication by enhancing antibody generation. This study underscores the importance of circulating Tfh cells in promoting non-neutralizing antibodies during HIV-1 infection.

## Introduction

A safe and effective prophylactic vaccine remains the most efficient way of ending the HIV/AIDS epidemic which affects over 36 million people worldwide (1). Although studies in non-human primate and animal models have demonstrated the efficacy of anti-HIV broadly neutralizing antibodies (bNAbs) in preventing HIV infection, human vaccine trials to date have been largely unsuccessful in inducing such responses (2–4). Thus, an improved understanding of the mechanisms that underlie the development of functional and durable anti-HIV antibody responses in the context of a natural infection will be essential for optimal vaccine design efforts (5). Moreover, with the quality of immune responses in early acute HIV infection predicting disease outcome (6, 7), early acute HIV infection is a useful model to identify early correlates of HIV-1 control.

T follicular helper (Tfh) cells, a lineage of CD4^+^ T cells that express the chemokine receptor CXCR5, are specialized for B cell help and the development of antibody responses (8, 9). Tfh-B cell interactions in the B cell follicles promote germinal center (GC) formation, B cell differentiation, B cell survival, antibody affinity maturation and class switch recombination (8, 10). The circulating memory counterparts of *bona fide* germinal center Tfh cells have been recently described (11, 12). These cells display either an activated or quiescent phenotype based on the expression of PD-1 and ICOS or CCR7 receptors and can be further divided into subsets based on the expression of CXCR3 and CCR6 receptors (12, 13). The subsets; Tfh1, Tfh2, Tfh17 and Tfh1-17, were named due to their similarities to other T helper cell lineages. Tfh1 cells express CXCR3 like Th1 cells, Tfh2 cells produce IL-4 like Th2 cells, Tfh17 cells express CCR6 similar to Th17 cells and Tfh1-17 cells have functional properties that are similar to both Th1 and Th17 cells (12–14).

From the RV144 vaccine trial, which had a modest efficacy in preventing HIV acquisition, we learned that non-neutralizing antibodies (nnAbs) could protect against HIV acquisition (15). Consistent with this observation, a recent study exploring the efficacy of nnAbs for blocking virus entry, showed that anti-Env nnAbs could modulate the transmission of simian HIV (SHIV) in macaques and reduce the number of transmitted/founder viruses establishing infection in the animals (16). Moreover, a humanized mouse model of HIV infection, reported near-complete clearance of adoptively transferred infected cells within 5 hours of nnAbs treatment (17) further demonstrating the potential for nnAbs in preventing HIV infection. Specific Tfh subsets have been shown to help the induction of various antibody functions. For instance, a recent study correlated the frequencies of CXCR3^−^ cTfh; which includes both Tfh2 and Tfh17 subsets, with the development of bNAbs against HIV infection (18), suggesting a potential role of these subsets as correlates for the induction of bNAbs in infection and possibly by vaccines. It is thus important to define specific Tfh subsets that contribute to nnAbs development in the context of natural HIV infection.

Here we investigated if the induction of cTfh responses during acute HIV infection contribute to initial HIV control and promote the development of anti-HIV nnAbs. We examined the role of HIV-specific cTfh cell subsets during acute HIV infection using HLA class II tetramers and multiparametric flow cytometry. HIV-specific antibody responses were further measured using a customized multivariate Luminex assay. Our results showed that acute HIV infection induces significant expansion of HIV-specific memory Tfh1 cells (p=0.02), which correlated with lower set point viral loads. Moreover, the frequencies of Tfh1 cells during early infection were predictive of p24-specific IgG titers. These data suggest that circulating Tfh1 cells play a role in controlling viral replication during primary HIV infection by enhancing robust anti-HIV antibody production, which is desirable for a prospective HIV vaccine.

## Results

### Circulating CXCR5^+^ cells in healthy donors have a predominantly central memory phenotype

Recent studies have focused on characterizing circulating CXCR5^+^CD4^+^ T cells (cTfh) because of their similarities with germinal center Tfh cells and their potential role in the development of bNAbs (18, 19). The difficulty associated with obtaining *bona fide* Tfh from lymphoid tissues has also stirred the interest in studying cTfh as surrogates. Although the phenotype of cTfh has not been clearly defined, the consensus is that they represent circulating memory Tfh (13). To determine how HIV infection perturbs global frequencies and phenotypes of peripheral Tfh we began by establishing baseline characteristics of this cell population in our study cohort who are predominantly of the Zulu/Xhosa ethnicity. We used CCR7 and CD45RA, well established memory markers to define four memory subsets. Specifically, we defined naïve (N) T cells by gating on CCR7^+^ and CD45RA^+^, central memory (CM) by CCR7^+^CD45RA^−^, effector memory (EM) by CCR7^−^CD45RA^−^, and terminally differentiated effector memory (TEMRA) by CCR7^−^CD45RA^+^ (20) (Figure 1A). Phenotypic analysis of total CD4^+^ T cells from 12 HIV negative donors revealed that 34.0% (29.1-43.2) were naïve, 21.8% (19.1-28.0) were CM, 33.7% (30.4-44.4) were EM and 2.8% (2.1-3.3) were TEMRA (Figure 1B). Next, we measured the frequency of cTfh (CXCR5^+^CD4^+^) cells and found that they comprised 12% (10.1-14.3) of circulating CD4^+^ T cells (Figure 1C). Memory phenotyping of Tfh cells showed that cTfh cells comprised 37.3% of CM CD4^+^ T cells, 7.8 % of EM CD4^+^ T cells and only a paltry 2.6% and 2.9% of the naïve and TEMRA CD4^+^ T cell compartments respectively (Figure 1D). Consistent with studies in Caucasian populations (21, 22), our data show that cTfh constitute a small fraction of circulating CD4^+^ T cells and are predominantly of a central memory phenotype.

**Figure 1:**
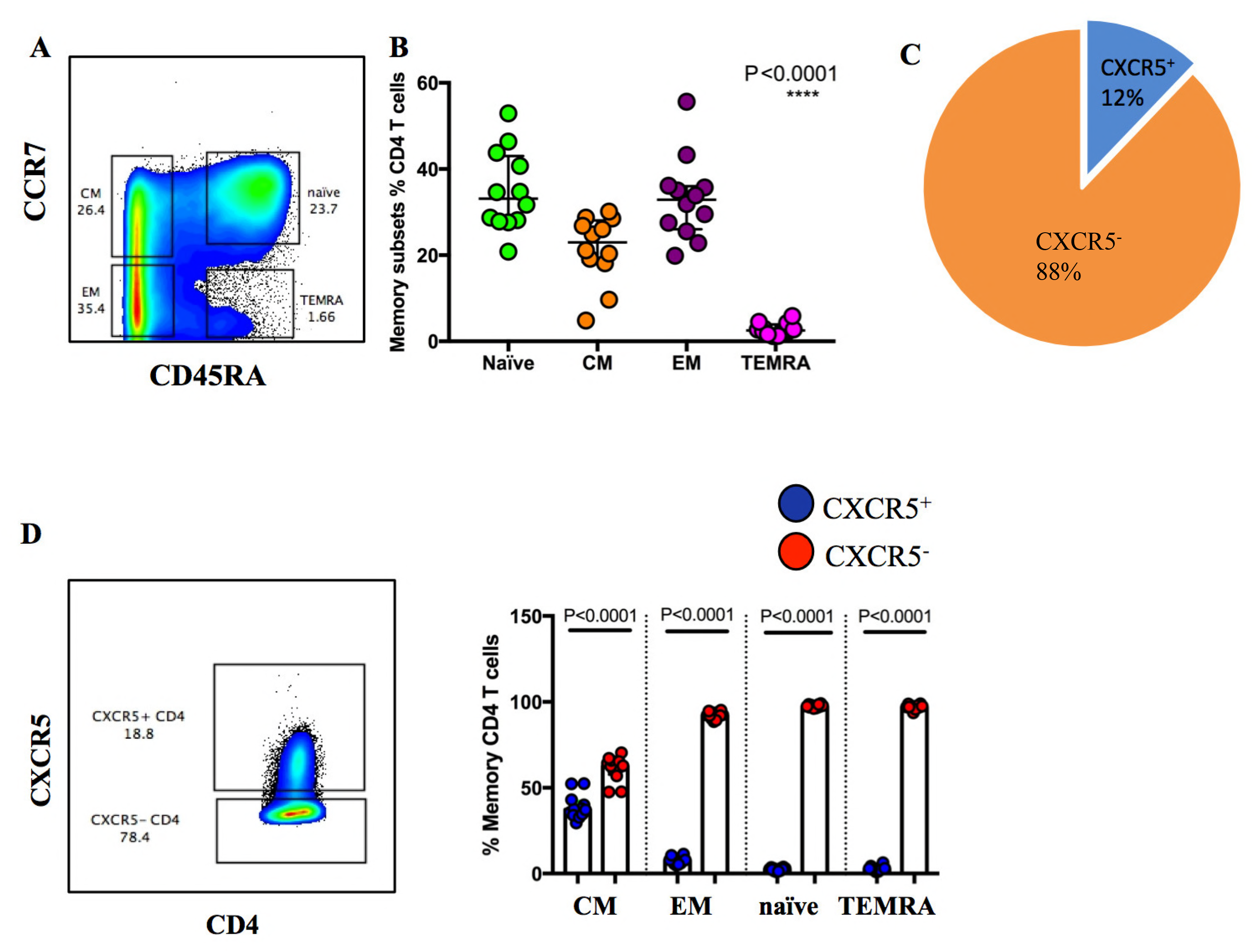
Memory distribution of CXCR5^+^ cells within circulating CD4^+^ T cell compartment in healthy donors. (A) Representative flow cytometry plot showing the gating strategy for CD4^+^ T cell memory populations. (B) Summary dot plots showing proportions of CD4^+^ T cells that are naïve, central (CM), effector (EM) and terminally differentiated (TEMRA) memory cells. (C) Pie chart showing median percentages of CXCR5^+^ and CXCR5^−^ CD4^+^ T cells. (D) Representative flow cytometry plots for CXCR5^+^ and CXCR5^−^ gating within bulk CD4^+^ T cells and summary plots depicting the proportions of CXCR5^+^ (blue) and CXCR5^−^ (red) CD4^+^ T cells within the CM, EM, naïve and TEMRA memory subsets. Statistical analysis was done using Kruskal-Wallis H test (B) and Mann-U Whitney tests (D).

### Perturbation of circulating Tfh cells during acute HIV-1 infection

Having established the normal frequencies and phenotypes of circulating Tfh cells, we next investigated how acute HIV infection alters the frequency and differentiation profiles of these cells. Samples obtained at a median of 6.9 weeks after HIV diagnosis were used for these studies (Table 1). As shown in (Figure 2A), HIV infection did not alter the overall frequencies of total circulating memory Tfh. However, memory subset analysis revealed an increase in naïve Tfh (p=0.004) and TEMRA Tfh (p=0.02), whereas CM Tfh (p=0.13) and EM Tfh (p=0.16) remained unchanged (Figure 2B).

**Table 1:** Characteristics of study participants

|  | **HIVneg** | **Acute HIV** |
| --- | --- | --- |
| **n** | 12 | 14 |
| **Male** | 0 | 6 |
| **Female** | 10 | 8 |
| **CD4 Count (cells/μl)\*** | N/A | 463 (422-561) |
| **Viral load (copies/ml)\*** | N/A | 121 000 (8 984-352 000) |
| **Time post-infection (weeks)\*** | N/A | 6.9 (5.0-8.0) |

\*Data are represented as median (IQR).

**Figure 2:**
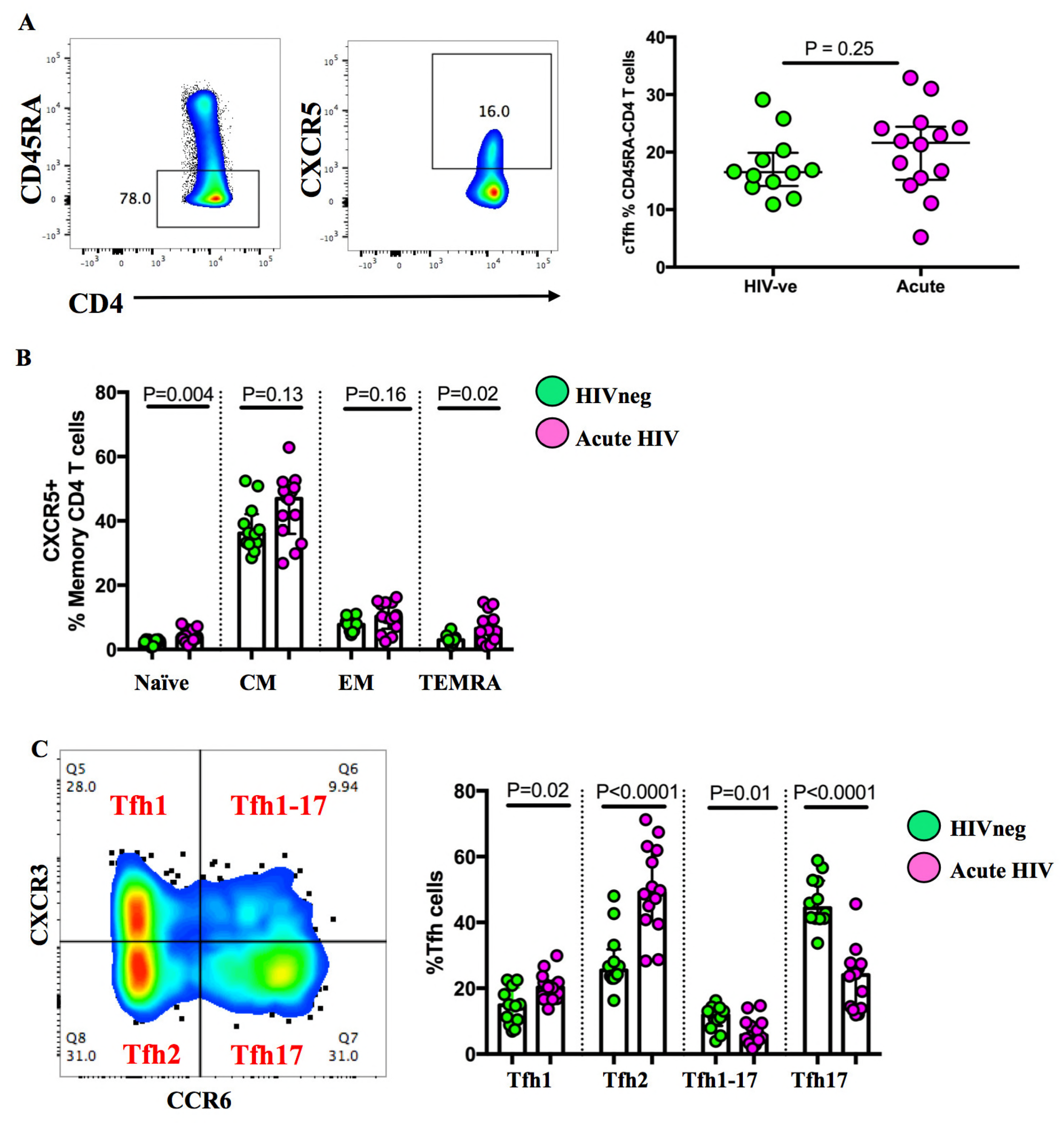
Heterogeneity within circulating Tfh compartment during acute HIV-1 infection. (A) Representative flow cytometry plots showing the gating strategy for bulk cTfh within CD45RA-CD4^+^ T cells and summary proportions of cTfh cells in HIV negative and acute HIV groups. (B) Summary plot comparing the frequencies of naïve, CM, EM and TEMRA cTfh cells in HIV negative and acute HIV donors. (C) Gating strategy for Tfh1, Tfh1-17, Tfh2 and Tfh17 subsets. The proportions of Tfh1, Tfh2, Tfh1-17 and Tfh17 subsets are compared between HIV negative and acute HIV groups. P values are from Mann-U Whitney tests.

Next, we used CXCR3 and CCR6 chemokine receptor markers to characterize cTfh subsets in an effort to identify which subset most influences the generation of anti-HIV antibodies during acute HIV infection. CXCR3 and CCR6 chemokine receptor markers have been previously used to identify several functional subsets that exhibit distinct B cell helper functions namely: Tfh1 (CXCR3^+^CCR6^−^), Tfh2 (CXCR3^−^CCR6^−^), Tfh1-17 (CXCR3^+^CCR6^+^) and Tfh17 (CXCR3^−^ CCR6^+^) (13). A representative flow plot, as seen in Figure 2C, depicts the distribution of cTfh subsets in an acutely infected donor based on the expression levels of the two respective chemokine receptor markers. Interestingly, acute infection skewed the distribution of cTfh subsets towards the Tfh1 (p=0.02) and Tfh2 (p<0.0001) phenotypes with significant reduction in the proportions of Tfh1-17 (p=0.01) and Tfh17 (p<0.0001) compared to HIV negative donors (Figure 2C).

### Frequency of Tfh1 cells during early acute HIV-1 infection correlates negatively with set point viral load

Having observed a significant expansion of Tfh1 and Tfh2, we next investigated if there was a relationship between the expanded cTfh subsets and set point viral load (SPVL), which is a reliable predictor of the rate of HIV disease progression. We calculated SPVL as the average VL from 3 to 12 months’ post-infection as previously reported (23) and correlated it to the frequencies of different cTfh subsets. Strikingly, Tfh1 frequencies correlated negatively with SPVL (p=0.03, r=−0.60) (Figure 3A) but there were no significant associations between Tfh2, Tfh1-17, Tfh17 or bulk cTfh cells and SPVL (Figures 3B, 3C, 3D and 3E). These results suggest that Tfh1 cells contribute to viral control during the first year of infection.

**Figure 3:**
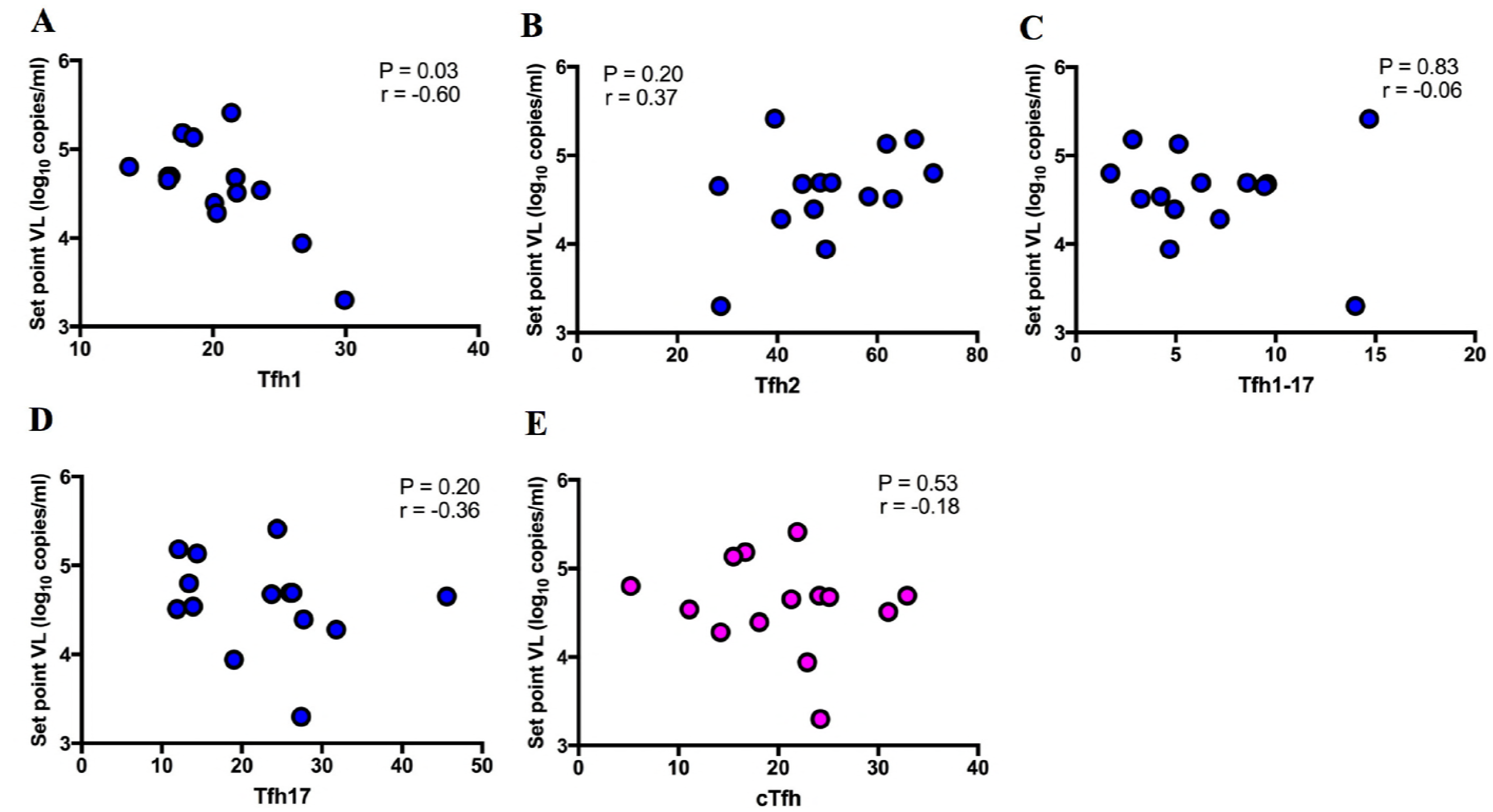
Tfh1 correlates negatively with set point viral load. Set point viral load was plotted against the frequency percentages of (A) Tfh1, (B) Tfh2, (C) Tfh1-17, (D) Tfh17 and (E) bulk cTfh cells determined by flow cytometry. Spearman rho (r) values and p values are reported.

### Tfh1 responses during early acute infection correlate with p24 IgG responses detected at one year post-infection

Numerous studies have associated slower disease progression with higher serum levels of HIV-1 Gag-specific IgG antibodies [reviewed in (24)]. We next, hypothesized that Tfh1 responses impact SPVL by driving the production of HIV-specific IgG antibodies. We measured plasma gp41, gp120, p17 and p24-specific IgG antibody titers at 12 months after infection for 10 study participants based on sample availability. Correlation analysis of IgG titers with SPVL revealed a negative correlation between SPVL and p24 IgG (p=0.007 r=−0.81) (Figure 4A) or gp41 IgG (p=0.009, r=−0.80) (Figure 4B) and no significant correlations between SPVL and p17 IgG (p=0.09, r=−0.58) (Figure 4C) or gp120 IgG titers (p=0.20, r=−0.44) (Figure 4D). We also examined the correlation of SPVL to the titers of p24 IgG isotypes; IgG1, IgG2, IgG3 and IgG4 and found no significant correlations between the p24 IgG isotypes and SPVL (Figure 4E and data not shown).

**Figure 4:**
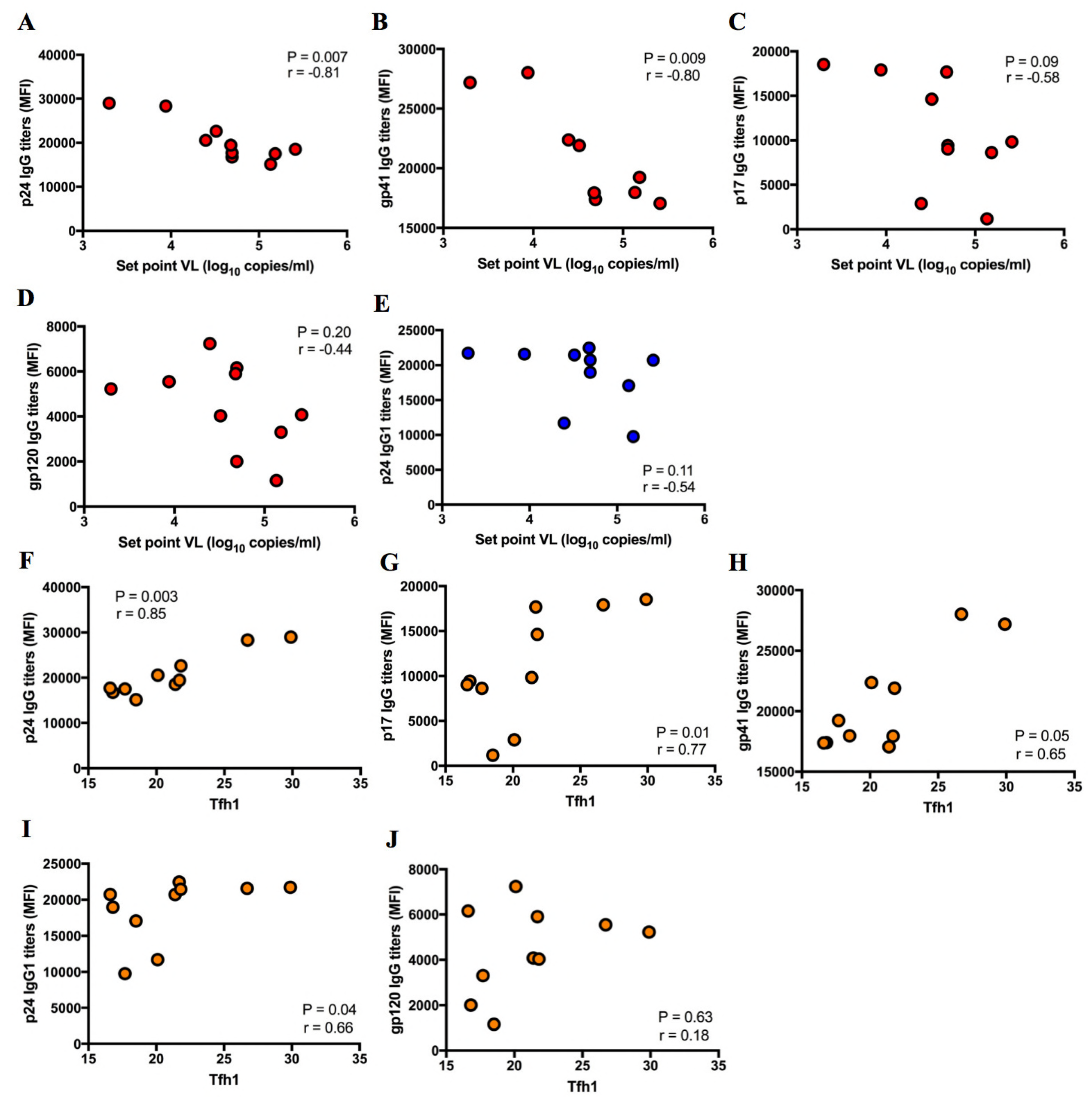
Early Tfh1 responses are predictive of p24 IgG responses at 1 year of infection. (A) p24 IgG titers, (B) gp41 IgG titers, (C) p17 IgG titers, (D) gp120 IgG titers and (E) p24 IgG1 titers, at 1 year time-point were determined using a customized multivariate Luminex assay and the values were inversely correlated to SPVL. (F) p24 IgG, (G) p17 IgG, (H) gp41 IgG, (I) p24 IgG1 and (J) gp120 IgG titers at 1 year time-point were correlated to the frequencies of Tfh1 at 6.9 (IQR, 5.0-8.0) weeks of infection. Mean Fluorescence Intensity (MFI) and viral load (VL). Spearman rho (r) values and p values are reported.

Lastly, we interrogated the relationship between Tfh1 frequencies and antibody titers. We found that Tfh1 frequencies during early infection (5.0-8.0 weeks) were directly correlated to the plasma titers of p24 IgG (p=0.003, r=0.85), p17 IgG (p=0.01, r=0.77), gp41 IgG (p=0.05, r=0.65) and p24 IgG1 (p=0.04, r=0.66) that were detected at 1 year post-infection (Figures 4F, 4G, 4H and 4I). There was however, no association between gp120 titers and Tfh1 frequencies (Figures 4J). These results suggest that the polarization of cTfh responses towards a Tfh1 phenotype can potentially impact the development of long-lasting antibody responses.

### HIV-specific Tfh responses are induced during acute HIV-1 infection

Next, we investigated if the expanded cTfh in acute HIV infection were HIV-specific using intracellular cytokine staining (ICS) assay and MHC class II tetramers. Although HIV-specific CD4^+^ T cells are important for viral control (25), the presence of HIV-specific Tfh responses in circulation remains controversial (11, 18). Therefore, we interrogated the cytokine expression pattern of cTfh cells after stimulation with HIV peptides. Figure 5A shows representative flow plots of unstimulated controls (top panel), cytokine secreting antigen specific CD4^+^ cells responding to HIV peptide pools (middle panel) or SEB stimulation (bottom panel) in an ICS assay. Our group previously showed that most of the HIV-specific CD4^+^ T cells in chronic clade C infection target the HIV Gag protein (26). Here we found no significant differences in Gag, Nef and Env responses (Figure 5B). Further interrogation of the cytokine profile of cTfh cells revealed that unlike SEB-specific cells which abundantly secreted TNF-α and IFN-γ (Figure 5C), HIV-specific cTfh were biased towards the secretion of Tfh functional cytokines: IL-21 and IL-4, with lower proportions of cTfh cells secreting TNF-α and IFN-γ (Figure 5D). Comparative analysis with non cTfh cells revealed that HIV-specific cTfh cells (blue) secreted more IL-21 (Figure 5E i, ii & iv) and IL-4 (Figure 5E ii & iii) whereas non cTfh cells (red) secreted significantly more IFN-γ (Figure 5E i & iv).

**Figure 5:**
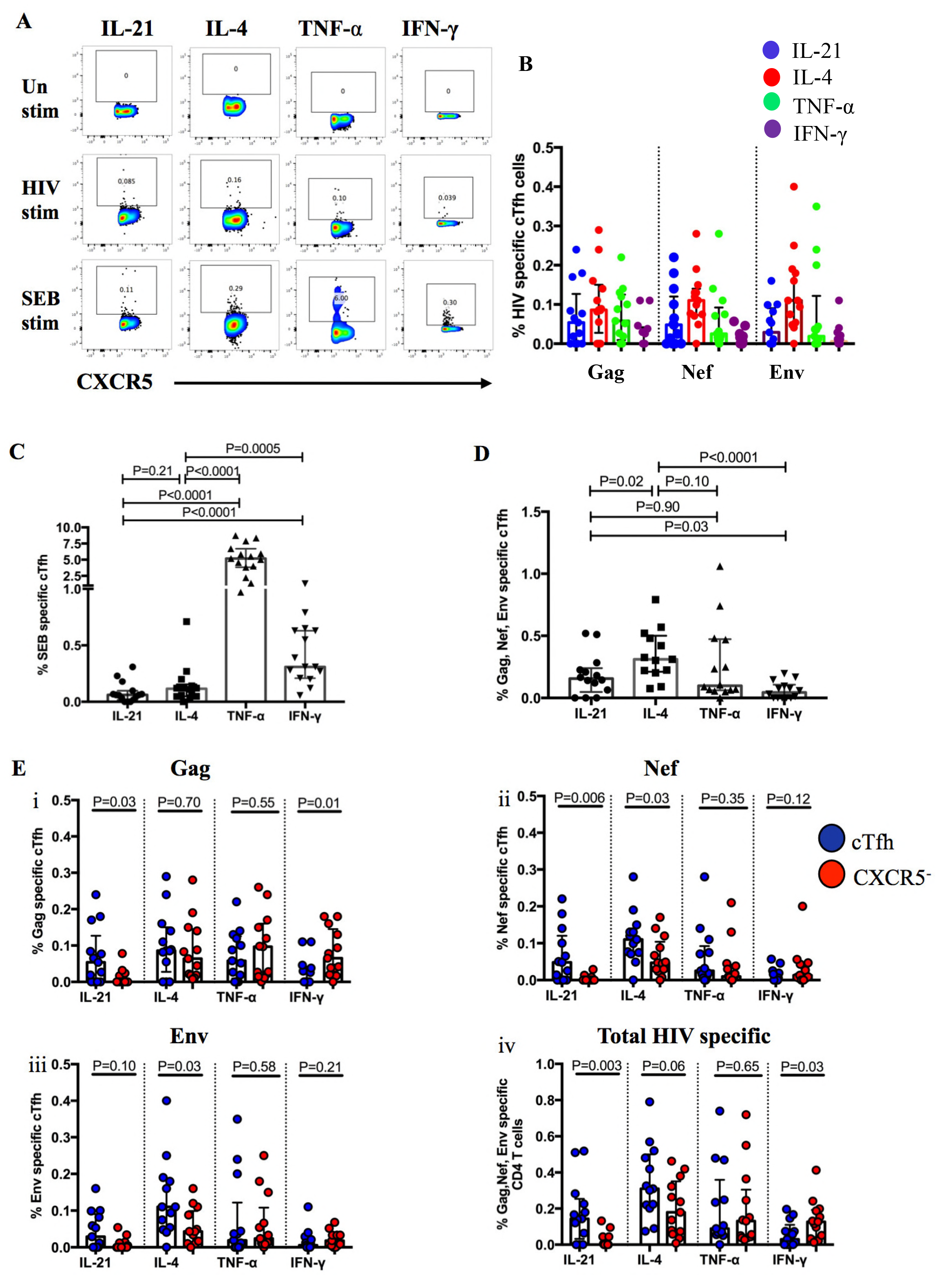
HIV specific cTfh measurements using ICS assay. (A) Representative flow cytometry plots for cytokine secreting cTfh cells. PBMCs were unstimulated or stimulated with SEB or HIV OLP pools for Gag, Nef and Env for 16h in the presence of GolgiStop and GolgiPlug transport inhibitors (BD Biosciences), and the intracellular expression of IL-21, IL-4, TNF-α and IFN-*γ* respectively was measured. (B) Summary frequency plots for Gag, Nef and Env-specific cTfh cells. (C) Summary plots for SEB stimulated cells. (D) Total HIV-specific cTfh cells. IL-21^+^, IL-4^+^, TNF-α^+^ and IFN-γ^+^ cTfh cells were summed up for Gag, Nef and Env. (E) Comparison of the cytokine secretion profiles of cTfh (CXCR5^+^) and non-cTfh (CXCR5^−^) cells. Frequencies for Gag (i), Nef (ii) and Env-specific (iii) cells were plotted separately or totaled (iv). P values are from Mann-U Whitney test.

### Persistence of Gag-specific Tfh responses during HIV-1 infection

We further used MHC class II tetramers to confirm the presence of HIV specific cTfh subsets and to track their dynamics over time. Using longitudinal samples for one acute HIV participant (patient 1) who had strong a response to the Gag C41 epitope restricted by DRB1*11:01 HLA haplotype, dual tetramer positive cells were detected within CD4^+^ T cells (Figure 6A). An overlay of these double tetramer CD4^+^ population onto CXCR5^+^CXCR3^+^CD4^+^ cTfh showed that HIV-specific cTfh were predominantly CXCR3^+^ (Tfh1 and Tfh1-17) cells and were detectable at 12, 14, 16 and 20 weeks post-infection (Figure 6B). Persistent HIV-specific cTfh cells during HIV-1 infection, was observed in another acute patient (patient 2) at 6 weeks and 138 weeks post infection (Figure 6C). Five additional patients expressing either the DRB1*11:01 (n=4) or the DRB1*13:01 (n=1) HLA haplotypes had detectable HIV-specific cTfh responses in chronic infection (>2 years of infection). Combined data for the 7 patients tested (Table 2) showed significantly higher frequencies of CXCR3^+^ (Tfh1 and Tfh1-17) tetramer-specific cells compared to Tfh2 and Tfh17 (p=0.0007) (Figure 6D). Together, these results demonstrate that HIV-specific cTfh cells persist during HIV infection.

**Table 2:** Study participants for tetramer staining assay

|  | **Acute HIV** | **Chronic HIV** |
| --- | --- | --- |
| **n** | 2 | 5 |
| **Male** | 0 | 0 |
| **Female** | 2 | 5 |
| **CD4 Count (cells/μl)\*** | 365 (351-433) | 720 (537-1 022) |
| **Viral load (copies/ml)\*** | 18 760 (9 642-377 905) | 2950 (455-30 975) |

\*Data are represented as median (IQR).

**Figure 6:**
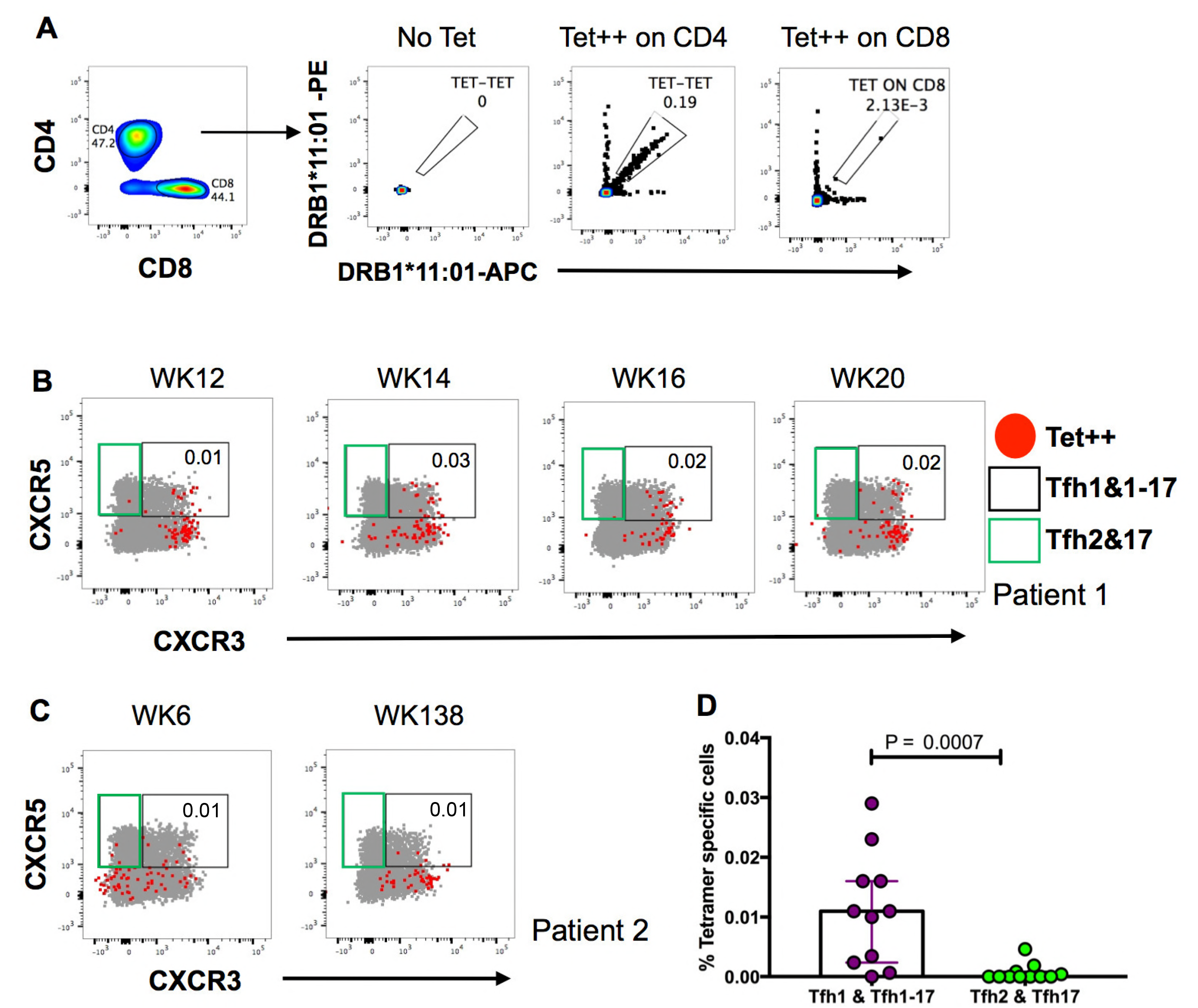
HIV specific cTfh detection by HLA class II tetramers. (A) Representative flow cytometry plots showing gating strategy for tetramer double positive (Tet++) cells within CD4^+^ T cells and CD8^+^ T cells. (B) HIV specific cTfh cells (in patient 1) are detected at 12, 14, 16 and 20 weeks (patient 1) or at (C) 6 and 138 weeks post-infection (patient 2) using HLA class II tetramers. Overlay of Tet++ cells (red) onto CXCR5^+^CXCR3^+^CD4^+^ cTfh. CXCR3^+^CXCR5^+^ gate (black) and CXCR3^−^CXCR5^+^ gate (green). (D) Summary plot comparing the frequencies of CXCR3^+^ (Tfh1 & Tfh1-17) and CXCR3^−^ (Tfh2 & Tfh17) tetramer specific cTfh cells. P value is from Mann-U Whitney test.

## Discussion

The extreme genetic diversity of HIV is a significant obstacle in the development of an effective anti-HIV vaccine (27). Even with the identification and isolation of several potent bNAbs in recent years, induction of such antibodies *in vivo* by vaccination has been a challenge (27, 28). Furthermore, nnAbs have been associated with protection from HIV acquisition and could be easier to induce by immunization as compared to bNAbs (29). This study sheds new light on circulating CD4^+^ T cell help that can impact the development of effective non-neutralizing anti-HIV antibody responses.

To understand how HIV modulates the frequency and function of circulating HIV-specific Tfh responses during primary HIV infection, we first established baseline frequencies of cTfh cells in HIV uninfected individuals. Comparative analysis between HIV infected and uninfected individuals showed there are similar frequencies of total memory cTfh cells across both groups. More in depth phenotypic characterization of cTfh cells revealed four distinct functional subsets namely Tfh1, Tfh2, Tfh1-17 and Tfh17 cells. We next showed that the increased frequency of Tfh1 cells positively correlated with p24 IgG antibody responses and negatively correlated with set point viral load. These data suggest that the Tfh1 subset plays an important role in the induction of anti-HIV antibodies and may contribute to control of HIV replication, consistent with murine model studies which have shown that cTfh cells can traffic into lymph nodes and interact with B cells in interfollicular zones and in germinal centers (30).

The differential induction of cTfh subsets has been described in the context of other infectious diseases. Consistent with our data, the early induction of circulating CXCR3^+^ cTfh, which comprises Tfh1 and Tfh1-17 subsets, correlated with the emergence of protective responses to the Influenza vaccine (31). In a subsequent study, they further demonstrated that CXCR3^+^ Tfh cells promote the development of high avidity antibody responses to the H1N1 vaccine (32). The aforementioned studies and our data suggest that Tfh1 cells might play an important helper role in the production of efficacious antiviral nnAbs. However, since studies using *in vitro* Tfh and B cell co-culture assays, have shown that CXCR3^+^ Tfh cells are effective in providing help to memory B cells but deficient at offering naïve B cell help (11, 18), more mechanistic work using animal models will be critical to delineating the intricacies of circulating Tfh1 cell helper capacity and providing clarity on the functional ability of Tfh1 subsets.

From our results, we also observed an expansion of the Tfh2 subsets during acute HIV-1 infection compared to the controls. The CXCR3^−^ subset which comprises Tfh2 and Tfh17 subsets has been described as having superior helper capacity *in-vitro* (11, 18) and the frequencies during acute HIV-1 infection was predictive of the ability to develop bNAbs in one study (18). However, another study did not see any relationship between this subset and the ability to develop bNAbs (22). Although, we sought to determine the relationship between the Tfh2 subset and bNAbs development in our study, only one study participant developed bNAbs thus, we were unable to make any conclusions.

Several reports have implicated bulk CD4^+^ T cells in immune mediated control of chronic HIV infection (25, 26, 33), but little is known about the role of HIV-specific cTfh cells in HIV control mainly because of their very low frequency in circulation and the paucity of reliable tools to study them. Even though there were few numbers of cytokine secreting cTfh cells in response to stimulation by HIV peptide as previously shown (34), our tetramer staining results provided conclusive evidence of the existence of HIV-specific cTfh cells during primary HIV infection. Notably, unlike bulk HIV-specific CD4^+^ T cells, which mostly target Gag, our data show that cTfh responses during acute HIV infection are dynamic and comprise a broad repertoire of cells specific for HIV-1 Gag, Nef and Env proteins. Virus-specific cTfh cells targeting different HIV proteins may have synergistic antiviral effect via cross-talk through the so-called intrastructural help to promote a greater net antiviral effect. This concept was first demonstrated in SIV_MAC_ Gag adenoviral vector immunized macaques and later validated by a murine model of SIV_MAC_ infection (35, 36). In the initial study, a faster onset and magnitude of antibody-dependent cell-mediated virus inhibition mediated by Env-specific antibodies, was observed in immunized animals compared to controls (35, 36). Human studies of cTfh cells comparing the effector profile of cTfh cells having different HIV-protein specificity, showed that Env-specific cTfh cells were superior at inducing class switching to IgG while Gag-specific cTfh cells were better at inducing B cell proliferation and maturation (37). These assays were conducted *in-vitro* but the micro-anatomy of immune responses *in-vivo* might encourage interactions between cells of different specificities. Additionally, studies have alluded to some degree of promiscuity in Tfh cell help to B cells in the GCs. It has been shown that the Tfh response is polyclonal (38), also the egression of Tfh cells from their initial colonized GCs and migration into other GCs have been documented (38, 39). These kinds of results argue for a less rigid Tfh help and highlight the dynamism of Tfh cell-B cell interactions which are the subject of many studies.

As previously mentioned, our tetramer staining results give a strong indication that cTfh cells persist in circulation well into chronic HIV infection. Although there were significantly higher frequencies of Tfh2 cells compared to Tfh1 cells during acute HIV, there was a higher proportion of tetramer specific Tfh1 cells. The tetramers we tested were directed at the Gag C41 epitope and one possibility is that Tfh2 cells may be targeting a different epitope other than the Gag C41 epitope which we interrogated. We however, consider the expansion of the Tfh2 subset as an interesting observation that warrants further studies.

Our data reveals an important association between Gag p24-specific nnAbs and viral load set point. The exact mechanism of how nnAbs influence HIV replication requires further investigation. Nevertheless, we speculate that the negative correlation between antibody titers and lower viral load set point may be attributable to antibody effector functions that have been associated with improved virus control (40) and slower HIV disease progression (41, 42). Although Fc effector functions like antibody-dependent cellular cytotoxicity (ADCC) and antibody-dependent cellular phagocytosis (ADCP) have been described as important for virus control, most of these functions were mostly attributed to Env-specific IgG antibodies. (41, 42). Studies investigating the mechanisms for virus suppression by Gag-specific antibodies have described the ability of these Gag-specific antibodies to opsonize antigens and recruit conventional or plasmacytoid dendritic cells to phagocytose antibody-coated antigens (43–45). Additionally, these opsonophagocytic IgG responses were associated with lower plasma HIV-RNA levels (43, 46), thus, highlighting this as the potential mechanism of virus control. Nevertheless, investigations into these and other antibody effector mechanisms are underway for this cohort.

Early studies investigating the kinetics and magnitude of anti-Gag and anti-Env IgG antibodies observed that the decay of Gag-specific antibodies correlated with poorer disease outcomes and argued that Gag-specific antibodies are a surrogate for CD4 T cell help to Gag-specific CD8 T cells (47). CD8 T cells are important for virus control and robust IL-21 mediated Tfh help to CD8 T cells improves CD8 T cell cytolytic activity (37), but we observed no correlations between the frequencies of IFN-γ^+^ CD8 T cells and lower set point viral loads among our study participants. Additionally, a paper from our group now in press showed that the association between Gag-p24 IgG and viral control was still maintained even after controlling for Gag-specific CD4 and CD8 T cell responses suggesting a CD8 T cell independent anti-viral mechanism of these antibodies (48).

A notable limitation of the study is the small sample size due to difficulty in recruiting subjects with untreated acute HIV-1 infection in the present era of mass ART induction in all HIV-1 infected patients. Nevertheless, despite the small sample size, we generated statistically significant results that provide new insight into the role of cTfh cells and their impact on the induction of antibody responses during primary HIV infection. Further studies to validate our findings in other acute infection cohorts are warranted.

In conclusion, the present study has identified a circulating Tfh1 subset whose frequency during acute HIV infection predicts the development of anti-p24 non-neutralizing antibodies. We also show that higher p24 IgG titers contribute to the control of HIV replication and have a beneficial effect on HIV disease progression. These results highlight the important role of HIV-specific cTfh cells in the generation of robust anti-HIV antibody responses, which are desirable for an HIV vaccine. Additionally, the identification of a cTfh subset that predicts the development of highly functional antibody responses might be useful to vaccine trials/studies as a potential biomarker to predict the development of robust antibody responses in vaccine responders or as a potential cell subset that can be manipulated to enhance vaccine responses (49).

## Materials and Methods

### Study Participants

Study participants comprised of 16 acute and 5 chronic HIV-infected ART-naïve individuals from HIV Pathogenesis Programme (HPP) Acute Infection cohort, Durban, South Africa. Patients were chosen based on availability of acute infection samples. Acute infection classification and disease staging in this cohort was previously described (23). Briefly, at screening, patients had detectable HIV RNA but had not yet seroconverted, either by ELISA or Western blotting. The date of infection for the study participants was estimated to be 14 days prior to screening as previously described (50). One acute infection time point was selected per patient for the study based on sample availability. The time post-infection across the patients was a median of 7 weeks (interquartile range-IQR, 5.25-7.75). The CD4 count, viral load and other patient characteristics are summarized in table 1. 10 HIV uninfected individuals from the Females Rising Through Education, Support and Health (FRESH) cohort (51, 52), also in Durban, South Africa were included as controls. The controls were chosen randomly based on sample availability at the time the study was conducted. The University of KwaZulu-Natal Biomedical Research Ethics Committee (BREC) and the Massachusetts General Hospital ethics review board approved the study. All study participants signed informed consent for participation in the study.

### Immunophenotyping

For surface phenotyping, frozen PBMCs were thawed, rested and stained using the LIVE/DEAD Aqua dead cell staining kit (Thermofisher scientific, Waltham MA, USA) as per manufacturer’s instructions, followed by staining with an antibody panel comprising: CD14 HV500 (BD Biosciences, San Jose, CA), CD19 HV500 (BD Biosciences), CD3 BV711 (BioLegend, San Diego, CA, USA), CD8 Qdot 800 (Life Technologies, Carlsbad, CA, USA), CD4 Qdot 655 (Life Technologies), CXCR5 AF488 (BD Biosciences), PD-1 BV421 (BioLegend), CCR6 PE (BioLegend), CXCR3 BV605 (BioLegend), CD45RA PE-Cy7 (BioLegend), CCR7 PerCp Cy5.5 (BioLegend) and CD27 APCH7 (BD Biosciences). For intracellular cytokine staining, peripheral blood mononuclear cells (PBMCs) were either left unstimulated or stimulated with HIV clade C overlapping peptide (OLP) pools spanning Gag, Nef, or Env proteins or staphylococcal enterotoxin B (SEB, 0.5 μg/ml) in the presence of GolgiStop and GolgiPlug protein transport inhibitors (BD Biosciences) for 16 hours at 37°C. Cells were surface stained, washed, fixed and permeabilized using the BD Cytofix/Cytoperm kit (BD Biosciences) according to manufacturer’s instructions. Cells were subsequently stained intracellularly with IL-2 PE (BD Biosciences), IL-21 APC (BioLegend), TNF-α A700 (BD Biosciences), IL-4 BV605 (BioLegend) and IFN-γ PE-Cy7 (BioLegend) antibodies. Cells were acquired using an LSRFortessa cytometer (BD Biosciences) with FACSDiva^™^ software and fluorescence minus one controls were used to define gates for the different cell subsets. Data was analysed using the FlowJo version 10.0.8 (Flowjo, LLC).

### HLA class II tetramer staining

HIV-specific cTfh responses were measured using HLA class II tetramers. The immunodominant Gag C41 epitope (26) was interrogated using DRB1*11:01 and DRB1*13:01 tetramers produced in the laboratory of Dr Søren Buus as previously described (53). The design and validation of these tetramers by our group have also been described (26). Briefly, recombinant human DRB1*11:01 or DRB1*13:01 HLA molecules were complexed with clade C HIV-1 Gag 41 peptide (YVDRFFKTLRAEQATQDV). For the assay, PBMCs were stained for 1 hour at 37°C with APC and PE conjugated HLA class II tetramer complexes, washed in 2% fetal calf serum (FCS) in phosphate buffered saline (PBS) and then stained with these antibodies: LIVE/DEAD Fixable Blue dead cell stain kit (Thermofisher Scientific), CD3 BV711 (BioLegend), CD4 BV650 (BD Biosciences), CD8 BV786 (BD Biosciences), CXCR5 AF488 (BD Biosciences), CXCR3 BV605 (BioLegend), PD-1 BV421 (BioLegend) and CD45RA AF700 (BioLegend); for 20 min at room temperature. Cells were washed and acquired on the LSRFortessa (BD Biosciences).

### Customized multivariate Luminex assay

Plasma HIV-1 specific antibodies were measured using a customized multivariate Luminex assay as previously described (54). Carboxylated fluorescent polystyrene beads (Biorad, Hercules, CA, USA) were coated with HIV-1 specific proteins including gp120 clade C of strain ZA.1197MB, gp41 clade C of strain ZA.1197MB, C-terminal 6xHis tagged p24 subtype C and p17 HXBc2 (Immune Technology, New York, NY, USA). Plasma samples were incubated with antigen-coated beads in a 96 well plate and unbound antibodies were washed with 0.05% Tween-20 in PBS. HIV-1 specific IgG antibodies were detected with phycoerythrin (PE) mouse IgG1-IgG4 secondary antibodies.

### Statistical analysis

Statistical analyses were performed using GraphPad Prism 7.0 (GraphPad Software, La Jolla, California, USA). Mann-Whitney U tests were used for the comparisons between any 2 groups. Variation across multiple groups was assessed using Kruskal-Wallis H test or two-way ANOVA. The correlation between two variables were done using Spearman’s rank correlation. P values were considered significant if less than 0.05.

## Acknowledgements

We would like to acknowledge all study participants and HPP Laboratory staff and thank Miss Fatima Laher (HPP, University of KwaZulu-Natal, South Africa) for assistance with HLA class II tetramer validation. We would also like to acknowledge the following funding sources; HHMI International research scholar award (Grant #55008743), The US National Institute of Health (R37 AI67073), Dan and Marjorie Sullivan Research scholar award (Grant # 224910), the support of the National Research Foundation (NRF) through a Doctoral Innovation scholarship to O.B. (2014-2016) and the South African Research Chairs Initiative. Additional funding was from the Mark and Lisa Schwartz Foundation, the Bill and Melinda Gates Foundation, the International AIDS Vaccine Initiative (IAVI), grant number UKZNRSA1001 and the Victor Daitz Foundation. This work was also partially supported by Gilead Sciences Incorporated and the Sub-Saharan African Network for TB/HIV Research Excellence (SANTHE), a DELTAS Africa Initiative (grant # DEL-15-006). The DELTAS Africa Initiative is an independent funding scheme of the African Academy of Sciences (AAS)’s Alliance for Accelerating Excellence in Science in Africa (AESA) and supported by the New Partnership for Africa’s Development Planning and Coordinating Agency (NEPAD Agency) with funding from the Wellcome Trust (grant # 107752/Z/15/Z) and the UK government. The views expressed in this publication are those of the author(s) and not necessarily those of AAS, NEPAD Agency, Wellcome Trust or the UK government.

## References

1. UNAIDS. 2017. Facts sheet.

2. Rerks-Ngarm S, Pitisuttithum P, Nitayaphan S, Kaewkungwal J, Chiu J, Paris R, Premsri N, Namwat C, de Souza M, Adams E, Benenson M, Gurunathan S, Tartaglia J, McNeil JG, Francis DP, Stablein D, Birx DL, Chunsuttiwat S, Khamboonruang C, Thongcharoen P, Robb ML, Michael NL, Kunasol P, Kim JH, Investigators M-T. 2009. Vaccination with ALVAC and AIDSVAX to prevent HIV-1 infection in Thailand. N Engl J Med 361:2209–2220.

3. Genesca M, Miller CJ. 2010. Use of nonhuman primate models to develop mucosal AIDS vaccines. Curr HIV/AIDS Rep 7:19–27.

4. Kwong PD, Mascola JR, Nabel GJ. 2011. Rational design of vaccines to elicit broadly neutralizing antibodies to HIV-1. Cold Spring Harb Perspect Med 1:a007278.

5. Martin-Gayo E, Cronin J, Hickman T, Ouyang Z, Lindqvist M, Kolb KE, Schulze Zur Wiesch J, Cubas R, Porichis F, Shalek AK, van Lunzen J, Haddad EK, Walker BD, Kaufmann DE, Lichterfeld M, Yu XG. 2017. Circulating CXCR5+CXCR3+PD-1lo Tfh-like cells in HIV-1 controllers with neutralizing antibody breadth. JCI Insight 2:e89574.

6. Pantaleo G, Demarest JF, Schacker T, Vaccarezza M, Cohen OJ, Daucher M, Graziosi C, Schnittman SS, Quinn TC, Shaw GM, Perrin L, Tambussi G, Lazzarin A, Sekaly RP, Soudeyns H, Corey L, Fauci AS. 1997. The qualitative nature of the primary immune response to HIV infection is a prognosticator of disease progression independent of the initial level of plasma viremia. Proc Natl Acad Sci U S A 94:254–258.

7. Deeks SG, Kitchen CM, Liu L, Guo H, Gascon R, Narvaez AB, Hunt P, Martin JN, Kahn JO, Levy J, McGrath MS, Hecht FM. 2004. Immune activation set point during early HIV infection predicts subsequent CD4+ T-cell changes independent of viral load. Blood 104:942–947.

8. Crotty S. 2011. Follicular helper CD4 T cells (TFH). Annual Review of Immunology 29:621–663.

9. Crotty S. 2014. T follicular helper cell differentiation, function, and roles in disease. Immunity 41:529–542.

10. Vinuesa CG, Tangye SG, Moser B, Mackay CR. 2005. Follicular B helper T cells in antibody responses and autoimmunity. Nat Rev Immunol 5:853–865.

11. Morita R, Schmitt N, Bentebibel S-E, Ranganathan R, Bourdery L, Zurawski G, Foucat E, Dullaers M, Oh S, Sabzghabaei N, Lavecchio EM, Punaro M, Pascual V, Banchereau J, Ueno H. 2011. Human Blood CXCR5(+)CD4(+) T Cells Are Counterparts of T Follicular Cells and Contain Specific Subsets that Differentially Support Antibody Secretion. Immunity 34:108–121.

12. Hale JS, Ahmed R. 2015. Memory T follicular helper CD4 T cells. Front Immunol 6:16.

13. Schmitt N, Bentebibel SE, Ueno H. 2014. Phenotype and functions of memory Tfh cells in human blood. Trends Immunol 35:436–442.

14. Morita R, Schmitt N, Bentebibel SE, Ranganathan R, Bourdery L, Zurawski G, Foucat E, Dullaers M, Oh S, Sabzghabaei N, Lavecchio EM, Punaro M, Pascual V, Banchereau J, Ueno H. 2011. Human blood CXCR5(+)CD4(+) T cells are counterparts of T follicular cells and contain specific subsets that differentially support antibody secretion. Immunity 34:108–121.

15. Haynes BF, Gilbert PB, McElrath MJ, Zolla-Pazner S, Tomaras GD, Alam SM, Evans DT, Montefiori DC, Karnasuta C, Sutthent R, Liao H-X, DeVico AL, Lewis GK, Williams C, Pinter A, Fong Y, Janes H, DeCamp A, Huang Y, Rao M, Billings E, Karasavvas N, Robb ML, Ngauy V, de Souza MS, Paris R, Ferrari G, Bailer RT, Soderberg KA, Andrews C, Berman PW, Frahm N, De Rosa SC, Alpert MD, Yates NL, Shen X, Koup RA, Pitisuttithum P, Kaewkungwal J, Nitayaphan S, Rerks-Ngarm S, Michael NL, Kim JH. 2012. Immune-Correlates Analysis of an HIV-1 Vaccine Efficacy Trial. New England Journal of Medicine 366:1275–1286.

16. Santra S, Tomaras GD, Warrier R, Nicely NI, Liao HX, Pollara J, Liu P, Alam SM, Zhang R, Cocklin SL, Shen X, Duffy R, Xia SM, Schutte RJ, Pemble Iv CW, Dennison SM, Li H, Chao A, Vidnovic K, Evans A, Klein K, Kumar A, Robinson J, Landucci G, Forthal DN, Montefiori DC, Kaewkungwal J, Nitayaphan S, Pitisuttithum P, Rerks-Ngarm S, Robb ML, Michael NL, Kim JH, Soderberg KA, Giorgi EE, Blair L, Korber BT, Moog C, Shattock RJ, Letvin NL, Schmitz JE, Moody MA, Gao F, Ferrari G, Shaw GM, Haynes BF. 2015. Human Non-neutralizing HIV-1 Envelope Monoclonal Antibodies Limit the Number of Founder Viruses during SHIV Mucosal Infection in Rhesus Macaques. PLoS Pathog 11:e1005042.

17. Horwitz JA, Bar-On Y, Lu CL, Fera D, Lockhart AAK, Lorenzi JCC, Nogueira L, Golijanin J, Scheid JF, Seaman MS, Gazumyan A, Zolla-Pazner S, Nussenzweig MC. 2017. Non-neutralizing Antibodies Alter the Course of HIV-1 Infection In Vivo. Cell 170:637–648 e610.

18. Locci M, Havenar-Daughton C, Landais E, Wu J, Kroenke MA, Arlehamn CL, Su LF, Cubas R, Davis MM, Sette A, Haddad EK, International AVIPCPI, Poignard P, Crotty S. 2013. Human Circulating PD-1(+)CXCR3(-)CXCR5(+) Memory Tfh Cells Are Highly Functional and Correlate with Broadly Neutralizing HIV Antibody Responses. Immunity 39:758–769.

19. Cohen K, Altfeld M, Alter G, Stamatatos L. 2014. Early Preservation of CXCR5+ PD-1+ Helper T Cells and B Cell Activation Predict the Breadth of Neutralizing Antibody Responses in Chronic HIV-1 Infection. Journal of Virology 88:13310–13321.

20. Larbi A, Fulop T. 2014. From “truly naïve” to “exhausted senescent” T cells: When markers predict functionality. Cytometry Part A 85:25–35.

21. Chevalier N, Jarrossay D, Ho E, Avery DT, Ma CS, Yu D, Sallusto F, Tangye SG, Mackay CR. 2011. CXCR5 expressing Human Central Memory CD4 T cells and their relevance for humoral immune responses. Journal of Immunology 186:5556–5568.

22. Boswell KL, Paris R, Boritz E, Ambrozak D, Yamamoto T, Darko S, Wloka K, Wheatley A, Narpala S, McDermott A, Roederer M, Haubrich R, Connors M, Ake J, Douek DC, Kim J, Petrovas C, Koup RA. 2014. Loss of circulating CD4 T cells with B cell helper function during chronic HIV infection. PLoS Pathog 10:e1003853.

23. Wright JK, Novitsky V, Brockman MA, Brumme ZL, Brumme CJ, Carlson JM, Heckerman D, Wang B, Losina E, Leshwedi M, van der Stok M, Maphumulo L, Mkhwanazi N, Chonco F, Goulder PJ, Essex M, Walker BD, Ndung’u T. 2011. Influence of Gag-protease-mediated replication capacity on disease progression in individuals recently infected with HIV-1 subtype C. J Virol 85:3996–4006.

24. French MA, Abudulai LN, Fernandez S. 2013. Isotype Diversification of IgG Antibodies to HIV Gag Proteins as a Therapeutic Vaccination Strategy for HIV Infection. Vaccines (Basel) 1:328–342.

25. Porichis F, Kaufmann DE. 2011. HIV-specific CD4 T cells and immune control of viral replication. Curr Opin HIV AIDS 6:174–180.

26. Laher F, Ranasinghe S, Porichis F, Mewalal N, Pretorius K, Ismail N, Buus S, Stryhn A, Carrington M, Walker BD, Ndung’u T, Ndhlovu ZM. 2017. HIV Controllers Exhibit Enhanced Frequencies of Major Histocompatibility Complex Class II Tetramer+ Gag-Specific CD4+ T Cells in Chronic Clade C HIV-1 Infection. J Virol 91.

27. Ahmed Y, Tian M, Gao Y. 2017. Development of an anti-HIV vaccine eliciting broadly neutralizing antibodies. AIDS Res Ther 14:50.

28. McCoy LE, Burton DR. 2017. Identification and specificity of broadly neutralizing antibodies against HIV. Immunol Rev 275:11–20.

29. Corey L, Gilbert PB, Tomaras GD, Haynes BF, Pantaleo G, Fauci AS. 2015. Immune correlates of vaccine protection against HIV-1 acquisition. Sci Transl Med 7:310rv317.

30. Sage PT, Alvarez D, Godec J, von Andrian UH, Sharpe AH. 2014. Circulating T follicular regulatory and helper cells have memory-like properties. J Clin Invest 124:5191–5204.

31. Bentebibel SE, Lopez S, Obermoser G, Schmitt N, Mueller C, Harrod C, Flano E, Mejias A, Albrecht RA, Blankenship D, Xu H, Pascual V, Banchereau J, Garcia- Sastre A, Palucka AK, Ramilo O, Ueno H. 2013. Induction of ICOS+CXCR3+CXCR5+ TH cells correlates with antibody responses to influenza vaccination. Sci Transl Med 5:176ra132.

32. Bentebibel SE, Khurana S, Schmitt N, Kurup P, Mueller C, Obermoser G, Palucka AK, Albrecht RA, Garcia-Sastre A, Golding H, Ueno H. 2016. ICOS(+)PD-1(+)CXCR3(+) T follicular helper cells contribute to the generation of high-avidity antibodies following influenza vaccination. Sci Rep 6:26494.

33. Schieffer M, Jessen H, Oster A, Pissani F, Soghoian DZ, Lu R, Jessen A, Zedlack C, Schultz B, Davis I, Ranasinghe S, Rosenberg ES, Alter G, Schumann R, Streeck H. 2014. Induction of Gag-specific CD4 T cell responses during acute HIV infection is associated with improved viral control. Journal of Virology 88:7357–7366.

34. Lindqvist M, van Lunzen J, Soghoian DZ, Kuhl BD, Ranasinghe S, Kranias G, Flanders MD, Cutler S, Yudanin N, Muller MI, Davis I, Farber D, Hartjen P, Haag F, Alter G, Schulze zur Wiesch J, Streeck H. 2012. Expansion of HIV-specific T follicular helper cells in chronic HIV infection. Journal of Clinical Investigation 122:3271–3280.

35. Liu J, O’Brien KL, Lynch DM, Simmons NL, La Porte A, Riggs AM, Abbink P, Coffey RT, Grandpre LE, Seaman MS, Landucci G, Forthal DN, Montefiori DC, Carville A, Mansfield KG, Havenga MJ, Pau MG, Goudsmit J, Barouch DH. 2009. Immune control of an SIV challenge by a T-cell-based vaccine in rhesus monkeys. Nature 457:87–91.

36. Nabi G, Genannt Bonsmann MS, Tenbusch M, Gardt O, Barouch DH, Temchura V, Uberla K. 2013. GagPol-specific CD4(+) T-cells increase the antibody response to Env by intrastructural help. Retrovirology 10:117.

37. Schultz BT, Teigler JE, Pissani F, Oster AF, Kranias G, Alter G, Marovich M, Eller MA, Dittmer U, Robb ML, Kim JH, Michael NL, Bolton D, Streeck H. 2016. Circulating HIV-Specific Interleukin-21(+)CD4(+) T Cells Represent Peripheral Tfh Cells with Antigen-Dependent Helper Functions. Immunity 44:167–178.

38. Vinuesa CG, Linterman MA, Yu D, MacLennan IC. 2016. Follicular Helper T Cells. Annu Rev Immunol 34:335–368.

39. Shulman Z, Gitlin AD, Targ S, Jankovic M, Pasqual G, Nussenzweig MC, Victora GD. 2013. T follicular helper cell dynamics in germinal centers. Science 341:673–677.

40. Ackerman ME, Mikhailova A, Brown EP, Dowell KG, Walker BD, Bailey-Kellogg C, Suscovich TJ, Alter G. 2016. Polyfunctional HIV-Specific Antibody Responses Are Associated with Spontaneous HIV Control. PLoS Pathog 12:e1005315.

41. Wren LH, Chung AW, Isitman G, Kelleher AD, Parsons MS, Amin J, Cooper DA, investigators Asc, Stratov I, Navis M, Kent SJ. 2013. Specific antibody-dependent cellular cytotoxicity responses associated with slow progression of HIV infection. Immunology 138:116–123.

42. Borrow P, Moody MA. 2017. Immunologic characteristics of HIV-infected individuals who make broadly neutralizing antibodies. Immunol Rev 275:62–78.

43. Tjiam MC, Taylor JP, Morshidi MA, Sariputra L, Burrows S, Martin JN, Deeks SG, Tan DB, Lee S, Fernandez S, French MA. 2015. Viremic HIV Controllers Exhibit High Plasmacytoid Dendritic Cell-Reactive Opsonophagocytic IgG Antibody Responses against HIV-1 p24 Associated with Greater Antibody Isotype Diversification. J Immunol 194:5320–5328.

44. French MA, Tjiam MC, Abudulai LN, Fernandez S. 2017. Antiviral Functions of Human Immunodeficiency Virus Type 1 (HIV-1)-Specific IgG Antibodies: Effects of Antiretroviral Therapy and Implications for Therapeutic HIV-1 Vaccine Design. Front Immunol 8:780.

45. Tjiam MC, Morshidi MA, Sariputra L, Martin JN, Deeks SG, Tan DBA, Lee S, Fernandez S, French MA. 2017. Association of HIV-1 Gag-Specific IgG Antibodies With Natural Control of HIV-1 Infection in Individuals Not Carrying HLA-B*57: 01 Is Only Observed in Viremic Controllers. J Acquir Immune Defic Syndr 76:e90–e92.

46. Tjiam MC, Sariputra L, Armitage JD, Taylor JP, Kelleher AD, Tan DB, Lee S, Fernandez S, French MA. 2016. Control of early HIV-1 infection associates with plasmacytoid dendritic cell-reactive opsonophagocytic IgG antibodies to HIV-1 p24. AIDS 30:2757–2765.

47. Binley JM, Klasse PJ, Cao Y, Jones I, Markowitz M, Ho DD, Moore JP. 1997. Differential regulation of the antibody responses to Gag and Env proteins of human immunodeficiency virus type 1. J Virol 71:2799–2809.

48. Chung A, Mabuka J, Ndlovu B, Licht A, Robinson H, Ramlakhan Y, Ghebremichael M, Reddy T, Goulder P, Walker B, Ndung’u T, Alter G. 2018. Viral Control in Chronic HIV-1 Subtype C Infection is associated with Enrichment of p24 IgG1 with Fc Effector Activity. AIDS In press.

49. Locci M, Havenar-Daughton C, Landais E, Wu J, Kroenke MA, Arlehamn CL, Su LF, Cubas R, Davis MM, Sette A, Haddad EK, Poignard P, Crotty S. 2013. Human circulating PD-(+)1CXCR3(-)CXCR5(+) memory Tfh cells are highly functional and correlate with broadly neutralizing HIV antibody responses. Immunity 39:758–769.

50. Van Loggerenberg F, Mlisana K, Williamson C, Auld SC, Morris L, Gray CM, Abdool Karim Q, Grobler A, Barnabas N, Iriogbe I, Abdool Karim SS, Team CAIS. 2008. Establishing a cohort at high risk of HIV infection in South Africa: challenges and experiences of the CAPRISA 002 acute infection study. PLoS One 3:e1954.

51. Ndhlovu ZM, Kamya P, Mewalal N, Kloverpris HN, Nkosi T, Pretorius K, Laher F, Ogunshola F, Chopera D, Shekhar K, Ghebremichael M, Ismail N, Moodley A, Malik A, Leslie A, Goulder PJ, Buus S, Chakraborty A, Dong K, Ndung’u T, Walker BD. 2015. Magnitude and Kinetics of CD8+ T Cell Activation during Hyperacute HIV Infection Impact Viral Set Point. Immunity 43:591–604.

52. Dong KL, Moodley A, Kwon DS, Ghebremichael MS, Dong M, Ismail N, Ndhlovu ZM, Mabuka JM, Muema DM, Pretorius K, Lin N, Walker BD, Ndung’u T. 2017. Detection and treatment of Fiebig stage I HIV-1 infection in young at-risk women in South Africa: a prospective cohort study. The Lancet HIV doi:https://doi.org/10.1016/S2352-3018(17)30146-7.

53. Braendstrup P, Justesen S, Osterbye T, Nielsen LL, Mallone R, Vindelov L, Stryhn A, Buus S. 2013. MHC class II tetramers made from isolated recombinant alpha and beta chains refolded with affinity-tagged peptides. PLoS One 8:e73648.

54. Brown EP, Licht AF, Dugast AS, Choi I, Bailey-Kellogg C, Alter G, Ackerman ME. 2012. High-throughput, multiplexed IgG subclassing of antigen-specific antibodies from clinical samples. J Immunol Methods 386:117–123.

